# On the Orientation of Entorhinal Grids

**DOI:** 10.1101/349373

**Authors:** Mikhail A. Lebedev, Alexei Ossadtchi

## Abstract

In the groundbreaking paper that eventually led to the 2014 Nobel prize in Physiology or Medicine, Hafting et al. (2005) reported that when rats forage for chocolate crumbs in a large open field, some neurons in their entorhinal cortex, called grid cells, exhibit crystalline-like responses to animal position, i.e. grids. Among several key findings documented in this article, the authors noted for the first time that the grids of different neurons can be tilted relative to each other, particularly if these neurons are far apart. In support of this claim, the researchers illustrated two neuronal subpopulations with a 7-10° difference in their grid orientations. Since these data are available online, we were able to reexamine these findings. Here we report several clarifications to the original observations of Hafting et al. First, we show that the relationship between the entorhinal grids is more complex than a single rotation: for the neuronal subpopulations analyzed by Hafting et al., one axis of the hexagonal grid is indeed tilted, but the other axes are not. Second, we show that local ensembles of entorhinal neurons are preferentially tuned to certain directions defined by the grid; this effect is unclear when single neurons are analyzed in isolation. Third, we argue that rat navigation traces are patterned instead of being random. For example, the orientation of the vector field representing average velocity appears to match the orientation of the neuronal grid. Overall, our observations indicate that additional insights into the function of entorhinal grids could be provided by ensemble-level analyses and thorough examination of the connection between the navigation behavior and neuronal patterns.

**Highlights:**

- While our examination of the online dataset from Hafting et al. generally confirms their original findings, several clarifications should be made.
- For the two neuronal subpopulations, where Hafting et al. reported a 7-10° relative tilt between the grids, only one of the grid axes is tilted, whereas the others are not.
- When spatial response fields are plotted for neuronal subpopulations instead of single neurons, it is clear that each subpopulation exhibits spatially periodic bands aligned with one of the grid axes.
- Navigation traces are not random and appear to match the orientation and periodicity of the neuronal grid.

## Introduction

The report of Hafting et al. is truly revolutionary (Hafting, Fyhn et al. 2005), as they clearly showed the existence of grid-like patterns in the positional responses of entorhinal neurons. Their previous report (Fyhn, Molden et al. 2004) was less convincing because of the smaller size of the behavioral arena. Currently, google scholar reports more than 2,200 citations of Hafting et al., which makes important any clarification of their original findings.

Hafting et al. extensively analyzed two particularly notable subpopulations of entorhinal neurons: one, called subpopulation 1 here, was recorded near the postrhinal border and the other, subpopulation 2, was recorded more ventrally in the opposite hemisphere. In their Fig. 2e, Hafting et al. showed that these subpopulations exhibited grid patterns that were rotated with respect to each other by 7-10° (i.e. approximately 15% of the principal grid angle). Since these data are shared online (https://www.ntnu.edu/kavli/research/grid-cell-data), we were able to reproduce this finding, but also discovered several new results.

**Figure 1.**
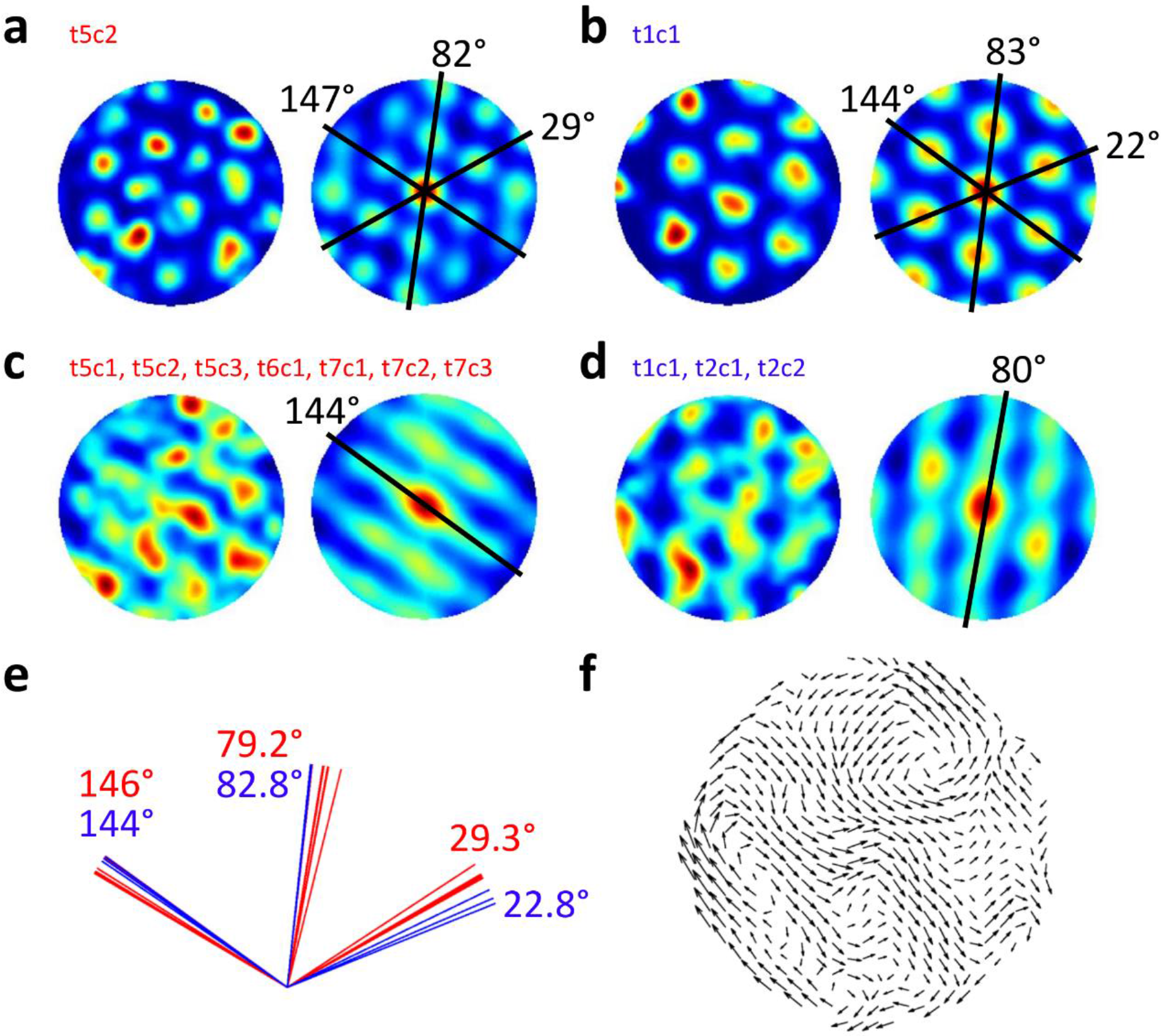
Analysis of neuronal grids. Data taken from the shared dataset: https://www.ntnu.edu/kavli/research/grid-cell-data. Original study: *Hafting*, *T*, *Fyhn*, *M*, *Molden*, *S*, *Moser*, *M-B*, *and Moser*, *EI (2005): Microstructure of a spatial map in the entorhinal cortex*. *Nature 436: 801-806*. **a**, Spatial map (left) and autocorrelation for a neuron from the first subpopulation. **b**, Spatial map (left) and autocorrelation for a neuron from the second subpopulation. **c**, Map and autocorrelation for all neurons from the first subpopulation lumped together. **d**, Map and autocorrelation for the second subpopulation. **e**, Orientation of grid axes for the first (red) and second (blue) subpopulations. Average angles are indicated for each subpopulation (color coded). **f**, Vector field for the average velocity of the rat in the arena.

## Results

Our analysis (Fig. 1) revealed the rotation reported by Haftin et al. only for one of the axes present in the grid: the one angled at 29.3° for subpopulation 1 and at 22.8° for subpopulation 2. The other two axes, at 80° and 145°, were virtually identical in both subpopulations, as evident from the spatial maps and autocorrelations for individual neurons (Fig. 1a,b) and the analysis of axes for all neurons (Fig. 1e).

Next, we assessed neuronal-population responses by lumping spikes from different neurons together for each subpopulation and calculating spatial maps and autocorrelations for these multiunit data (Fig. 1c,d). Our findings turned out to be different from the ones reported by Hafting et al. In their Fig. 3a, no interesting pattern could be discerned in the superimposed grids of the neurons from subpopulation 2. Hafting et al. commented that “grids from a small number of units recorded simultaneously at the same electrode position filled up the entire space of the recording arena.” Our population analysis revealed that the grid vertices did not fill the entire space but instead filled specific parts of the space: neurons from subpopulation 1 formed a pattern composed of bands parallel to the 145° axis, whereas the bands for subpopulation 2 were parallel to the 80° axis. Thus, not only the neuronal subpopulations recorded in different hemispheres shared the 80° and 145° axes of their grids (while exhibiting variability for the 20-30° axis), but also the grid vertices (from specific subpopulations) filled the paths defined by these axes.

Although Hafting et. al did not examine rat navigation traces in detail (and no such analysis can be found in the other papers from the same group), the orientation of the entorhinal grids could be related to the way the rats scanned the behavioral arena (Thompson, Berkowitz et al. 2017, Lebedev, Pimashkin et al. 2018). To this end, Fig. 1f shows the vector field of the average rat velocity. Patchy patterns that resemble the ones shown in Fig. 1a-d can be discerned. In particular, a vector flow can be seen in the middle of the plot with the orientation matching the 145° axis of the neuronal grid.

## Discussion

Based on our examination Hafting et al. dataset, we made three clarifications:

1. For the subpopulations of grid cells, where they reported a 7-10° tilt of the grid, such a tilt is present only for one of the axes (Fig. 1 a,b,e).
2. When spatial maps are plotted for local neuronal ensembles instead of single neurons, periodic bands are revealed oriented along one of the grid axes (Fig. 1c,d).
3. Vector field of navigation velocity contains a patchy pattern with the periodicity and orientation found in the neuronal grid (Fig. 1f).

Since we analyzed only 10 neurons available in the online database, our observations are somewhat anecdotal, so more data will have to be examined to reach a better understanding of the properties of local entorhinal ensembles. Yet, we find it intriguing that the signals generated by the neuronal ensembles from different hemispheres were “orthogonal” to each other (Kaufman, Churchland et al. 2014, Lebedev 2017), in the sense that their spatial receptive fields were aligned to different grid axes (Fig. 1c vs Fig. 1d).

Since the 2005 publication by Hafting et al., the Mosers group conducted additional investigations into the orientation of entorhinal grids (Stensola, Stensola et al. 2012, Stensola, Stensola et al. 2015). In one study (Stensola, Stensola et al. 2012), they conducted an analysis of several grid axes, instead of just one, and concluded that “grid cells cluster into a small number of layer-spanning anatomically overlapping modules with distinct scale, orientation, asymmetry and theta-frequency modulation”. While this conclusion is somewhat like our observations reported here, Stensola et al. did not focus on the possibility that different neuronal ensembles (or “modules” in their terminology) could share some of the grid axes. They also did not plot population spatial maps and did not reveal the bands aligned to the grid axes. (But neurons with such band-like spatial fields were reported by the O’Keefe group (Krupic, Burgess et al. 2012).) Another publication from the Mosers group (Stensola, Stensola et al. 2015) went further and suggested that “the systematic relationship between rotation and distortion of the grid pattern points to shear forces arising from anchoring to specific geometric reference points as key elements of the mechanism for alignment of grid patterns to the external world”. While this sounds like an interesting interpretation, it treats the grid is treated as a single entity that could be rotated or sheared and the possibility is not considered that grid axes instead of the grids themselves could be functionally significant.

While the researchers of the hippocampal formation are interested in how the geometry of the environment could shape the entorhinal grids (Stensola, Stensola et al. 2015, Krupic, Bauza et al. 2018), their research is not focused on the relationship between the environment geometry and navigation behavior, and between the navigation behavior and responses of entorhinal neurons. Recently, we (Lebedev, Pimashkin et al. 2018) and others (Thompson, Berkowitz et al. 2017) argued that the future studies of grid cells and place cells should pay more attention to different behaviors involved in navigations, including not just locomotion but also sniffing that could have a spatial function (Jacobs 2012, Corcoran, Pezzulo et al. 2018, Lebedev and Ossadtchi 2018, Tort, Brankačk et al. 2018). In addition to these arguments, our present report suggests that neuronal-ensemble properties should be considered, as well.

## Acknowledgement

This work was supported by the Center for Bioelectric Interfaces of the Institute for Cognitive Neuroscience of the National Research University Higher School of Economics, RF Government grant, ag. No. 14.641.31.0003.

## References

Corcoran, A. W., G. Pezzulo and J. Hohwy (2018). “Commentary: Respiration-Entrained Brain Rhythms Are Global but Often Overlooked.“ Frontiers in Systems Neuroscience 12(25).

Fyhn, M., S. Molden, M. P. Witter, E. I. Moser and M.-B., Moser (2004). “Spatial representation in the entorhinal cortex“. Science 305(5688): 1258–1264.

Hafting, T., M., Fyhn, S., Molden, M.-B., Moser and E. I. Moser (2005). “Microstructure of a spatial map in the entorhinal cortex“. Nature 436(7052): 801–806.

Jacobs, L. F. (2012). “From chemotaxis to the cognitive map: the function of olfaction“. Proceedings of the National Academy of Sciences 109(Supplement 1): 10693–10700.

Kaufman, M. T., M. M. Churchland, S. I. Ryu and K. V. Shenoy (2014). “Cortical activity in the null space: permitting preparation without movement“. Nature neuroscience 17(3): 440.

Krupic, J., M., Bauza, S., Burton and J., O’Keefe (2018). “Local transformations of the hippocampal cognitive map“. Science 359(6380): 1143–1146.

Krupic, J., N., Burgess and J., O’Keefe (2012). “Neural representations of location composed of spatially periodic bands“. Science 337(6096): 853–857.

Lebedev, M. (2017). “Commentary: Cortical activity in the null space: permitting preparation without movement“. Frontiers in Neuroscience 11: 502.

Lebedev, M. and A., Ossadtchi (2018). “Commentary: Spatial olfactory learning contributes to place field formation in the hippocampus“. Frontiers in Systems Neuroscience 12: 8.

Lebedev, M., A., Pimashkin and A., Ossadtchi (2018). “Navigation patterns and scent marking: underappreciated contributors to hippocampal and entorhinal spatial representations?“ Frontiers in Behavioral Neuroscience 12: 98.

Stensola, H., T., Stensola, T., Solstad, K., Frøland, M.-B., Moser and E. I. Moser (2012). “The entorhinal grid map is discretized“. Nature 492(7427): 72.

Stensola, T., H., Stensola, M.-B., Moser and E. I. Moser (2015). “Shearing-induced asymmetry in entorhinal grid cells“ Nature 518(7538): 207.

Thompson, S. M., L. E. Berkowitz and B. J. Clark (2017). “Behavioral and neural subsystems of rodent exploration“. Learning and Motivation.

Tort, A. B., J. Brankačk and A., Draguhn (2018). “Respiration-entrained brain rhythms are global but often overlooked“. Trends in neurosciences.

